# Absence of receptor for advanced glycation end product (RAGE) reduces inflammation and extends survival in the hSOD1^G93A^ mouse model of amyotrophic lateral sclerosis

**DOI:** 10.1101/2020.05.01.071944

**Authors:** John D Lee, Tanya S McDonald, Jenny NT Fung, Trent M Woodruff

## Abstract

Amyotrophic lateral sclerosis (ALS) is a fatal and rapidly progressing motor neuron degenerative disease that is without effective treatment. The receptor for advanced glycation end products (RAGE) is a major component of the innate immune system that has been implicated in ALS pathogenesis. However, the contribution of RAGE signaling to the neuroinflammation that underlies ALS neurodegeneration remains unknown. The present study therefore generated SOD1^G93A^ mice lacking RAGE, and compared them to SOD1^G93A^ transgenic ALS mice in respect to disease progression (i.e. body weight, survival and muscle strength), neuroinflammation and denervation markers in the spinal cord and tibialis anterior muscle. We found that complete absence of RAGE signaling exerted a protective effect on SOD1^G93A^ pathology, slowing disease progression and significantly extending survival by ∼3 weeks, and improving motor function (rotarod and grip strength). This was associated with reduced microgliosis, cytokines, innate immune factors (complement, TLRs, inflammasomes), and oxidative stress in the spinal cord, and a reduction of denervation markers in the tibialis anterior muscle. We also documented that RAGE mRNA expression was significantly increased in the spinal cord and muscles of preclinical SOD1 and TDP43 models of ALS, supporting a widespread involvement for RAGE in ALS pathology. In summary, our results indicate that RAGE signalling drives neuroinflammation and contributes to neurodegeneration in ALS, and highlights RAGE as a potential immune therapeutic target for ALS.

## INTRODUCTION

Amyotrophic lateral sclerosis (ALS) is a progressive heterogeneous neurodegenerative disease. It is characterized by the degeneration of the upper motor neurons in the motor cortex and lower motor neurons in the brainstem and spinal cord leading to muscle denervation, and atrophy, resulting in paralysis and eventual death due to respiratory muscle failure. The mechanisms leading to motor neuron death in ALS are still unclear, but an emerging body of evidence suggests that immune and inflammatory factors could contribute to the progression of the disease [1-3].

The receptor for advanced glycation end product (RAGE) is a member of the immunoglobulin superfamily and a cell surface receptor that plays a crucial role in inflammation, oxidative stress and cellular dysfunction in a number of neurodegenerative diseases [4]. RAGE is generally considered a pro-inflammatory pattern recognition receptor expressed by numerous immune and non-immune cells, including cells within the central nervous system [5,6,4]. It interacts with a number of molecules such as advanced glycation end products (AGE), S100/calgranulin family members, high mobility group box 1 (HMGB1), and molecules that are prone to aggregation, such as superoxide dismutase 1 (SOD1) and TAR DNA binding protein 43 (TDP-43) found in ALS [6,4]. This interaction triggers multiple intracellular signaling pathways that leads to the production of pro-inflammatory molecules and reactive oxygen species leading to oxidative stress and subsequent neuronal loss [7,8].

RAGE is also one of the major components of the innate immune system, which has been implicated as contributing to ALS pathology [9-12,6,13]. Multiple studies have demonstrated that RAGE and its inflammatory ligands AGE, S100β and HMGB1 are up-regulated in the spinal cord of human ALS patients compared to healthy controls [6,11]. Prior studies have also demonstrated that inhibition of RAGE using soluble RAGE form has beneficial effects on survival and disease progression in animal models of ALS [10,12], suggesting RAGE could play a pathogenic role in the disease. Although these studies have indicated detrimental actions of RAGE activation in ALS pathogenesis, the specific molecular mechanism by which RAGE contributes to neurodegeneration remains unknown.

In the present study, we demonstrated that RAGE is upregulated in degenerating tissues across multiple preclinical mouse models of ALS suggesting a common pathway to activation. We also identified the contribution of RAGE signaling in ALS pathogenesis by generating SOD1^G93A^ mice lacking RAGE and comparing them to SOD1^G93A^ mice in respect to survival, muscle strength and inflammatory status in the spinal cord and tibialis anterior (TA) muscle. Our findings demonstrate that lack of RAGE signaling has a protective effect on SOD1^G93A^ pathology, significantly extending survival and improving motor function. This was associated with reduced inflammation and oxidative stress in the spinal cord, and a reduction of denervation markers in the TA muscle. These results highlight RAGE as a potential immune therapeutic target for ALS.

## METHODS

### Animals

Transgenic SOD1^G93A^ mice (B6-Cg-Tg (SOD1-G93A) 1Gur/J) were obtained from the Jackson laboratory (Bar Harbor, ME, USA) and were bred on C57BL/6J background to produce SOD1^G93A^ and wild-type (WT) mice. Female SOD1^G93A^ and WT mice at four predefined stages of ALS were used in this study as described previously [14]. Specifically the predefined stages of disease progression are: 1) *pre-symptomatic* at 30 days postnatal where no motor deficits are seen; 2) *onset* at 70 days postnatal where there is initial signs of motor deficits determined by a significant reduction in hind-limb grip strength; 3) *mid-symptomatic* at 130 days postnatal where there is marked weakness in hind-limbs and tremor when suspended by the tail; 4) *end-stage* at 150 to 175 days postnatal where there is full paralysis of lower limbs and loss of righting reflex (also defined as the survival end-point). Transgenic TDP-43^WT^ (Line 96) and TDP-43^Q331K^ (Line 103) mice were obtained from the Jackson Laboratory (Bar Harbor, ME, USA) and were bred on a C57BL/6J background to produce TDP-43^WT^, TDP-43^Q331K^ and respective WT control mice. TDP-43^WT^ transgenic mice express a myc-tagged human non-mutated version of the TDP-43 cDNA sequence and TDP-43^Q331K^ mice express a myc-tagged, human TDP-43 cDNA modified to have an ALS-linked glutamine to lysine residue mutation at position 331, under the direction of the mouse prion protein promoter. The prion protein promoter ensures that the transgene expression is directed primarily to the central nervous system - the brain and spinal cord - and is very low in other peripheral tissue [15]. Female WT and TDP-43^Q331K^ mice at 16 months based on their motor deficits were used in this study [16,17]. Compound TDP-43^WTxQ331K^ mice were generated by breeding TDP-43^WT^ and TDP-43^Q331K^ mice as described previously [18]. Co-expression of WT and Q331K mutant form of TDP-43 resulted in a very aggressive motor phenotype and tremor. Female WT and TDP-43^WTxQ331K^ mice at 4 – 5 weeks were used due to the loss of righting reflex. RAGE^-/-^ female mice on C57BL/6 background were bred with male SOD1^G93A^ to yield SOD1^G93A^ mice lacking RAGE (SOD1^G93A^ x RAGE^-/-^) at F2 generation. Female SOD1^G93A^ and SOD1^G93A^ x RAGE^-/-^ mice were used for all phenotype studies. The breeding colony for all the animals was maintained at the University of Queensland Biological Resources Animal Facilities under specific pathogen free conditions. Animals were group housed (2-3 animals per cage) under identical conditions in a 12h light/dark cycle (lights on at 0600 h) with free access to food and water.

### Disease progression and survival analysis

The rate of disease progression in SOD1^G93A^ and SOD1^G93A^ x RAGE^-/-^ mice was determined by the age at which maximal grip strength declined by 25, 50, 75 and 100%. Survival was determined by the inability of the animal to right itself within 15 – 30 seconds if laid on either side. This is a widely accepted and published endpoint for life span studies in ALS mice [19] and guarantees that euthanasia occurs prior to the mice being unable to reach food or water.

### Evaluation of motor function and health

WT, SOD1^G93A^, RAGE^-/-^ and SOD1^G93A^ x RAGE^-/-^ mice (*n* = 15) were weighed weekly at the same time of the day (1600 h), from 42 days of age until the defined end stage (loss of righting reflex). A digital force gauge (Ugo Basile, Italy) was used to measure maximal hind-limb muscle grip strength as described previously. Mice were held by their tail and lowered until their hind-limbs grasped the T-bar connected to the digital force gauge. The tail was then lowered until the body was horizontal with the apparatus, and the mice were pulled away from the T-bar with a smooth steady motion until both of their hind-limbs released the bar. The strength of the grip was measured in gram force. Each mouse was given 10 attempts and the maximum grip strength from these attempts were recorded. Mice were also tested for their motor coordination using a Rota-rod apparatus (Ugo Basile, Italy) at a constant speed of 20 rpm. Each mouse was given three attempts and the longest latency to fall was recorded; 180s was chosen as the arbitrary cut off time. One week prior to the test, mice were trained twice to remain on the Rota-rod apparatus to exclude differences in motivation and motor learning. In the training phase, mice were placed on the Rota-rod at a constant speed of 20 rpm for a maximum duration of 240s [14].

### Real-time quantitative PCR

Total RNA was isolated from spinal cord and TA muscle of WT, SOD1^G93A^, RAGE^-/-^ and SOD1^G93A^ x RAGE^-/-^ mice (*n* = 5 - 6) using RNeasy Lipid Tissue extraction kit according to manufacturer’s instructions (QIAGEN, CA, USA). The total RNA was purified from genomic DNA contamination using Turbo DNase treatment (Ambion, NY, USA), then converted to cDNA using AffinityScript cDNA synthesis kit according to manufacturer’s instructions (Agilent Technologies, CA, USA). Commercially available gene-specific TaqMan probes for Ager (Mm 01134790_g1), Itgam (Mm00434455_m1), Cd68 (Mm03047343_m1), Aif1 (Mm00479862_g1), Gfap (Mm01253033_m1), Tnf (Mm00443258_m1), Il1b (Mm00434228_m1), C1qb (Mm01179619_m1), C3 (Mm01232779_m1), C5ar1 (Mm00500292_s1), Hmgb1 (Mm00849805_gH), Tlr4 (Mm00445273_m1), Nlrp3 (Mm00840904_m1), Pycard (Mm00445747_g1), Cybb (Mm01287743_m1), Musk (Mm01346929_m1), Chrna1 (Mm00431629_m1), Cpla2 (Mm00447040_m1), Ncam1 (Mm01149710_m1) and Runx1 (Mm01213404_m1) were used to amplify target gene of interest (Applied Biosystems, MA, USA). Relative target gene expression to geometric mean of reference genes Gapdh (Mm99999915_g1) and Actb (Mm02619580_g1) was determined using this formula: 2^-ΔCT^ where ΔCT = (Ct _(Target gene)_ – Ct _(Gapdh and Actb)_). Final measures are presented as relative levels of gene expression in SOD1^G93A^, TDP43^Q331K^ and TDP43^WTxQ331K^ mice compared with expression in WT mice and relative gene expression in. SOD1^G93A^, RAGE^-/-^ and SOD1^G93A^ x RAGE^-/-^ mice compared with expression in WT controls. Probe set was tested over a serial cDNA concentration for amplification efficiency. No reverse transcription and water as no template control were used as negative controls. All samples were run in triplicate and were tested in three separate experiments.

### Tissue preparation for microglia/astrocyte quantification and immunohistochemistry

SOD1^G93A^ and SOD1^G93A^ x RAGE^-/-^ mice at mid-symptomatic stage (*n* = 3) were euthanized by intraperitoneal injection of zolazapam (50 mg/kg; Zoletil, Lyppard) and xylazine (10 mg/kg; Xylazil, Lyppard). Mice were then fixed by transcardiac perfusion with 2% sodium nitrite in 0.1 M phosphate buffer (pH 7.4; Sigma-Aldrich, St Louis, MO, USA) followed by 4% paraformaldehyde in 0.1 M phosphate buffer (4% PFA-PB; pH 7.4; Sigma-Aldrich, St Louis, MO, USA). Lumbar spinal cords were collected and placed into 4% PFA-PB for 2 hours at 4 °C. Following this incubation, spinal cords were washed 3 x 5 min in phosphate buffered saline (PBS; pH 7.4), followed by submersion in sucrose solution at 15% then 30% in PBS (pH 7.4). Lumbar spinal cords were then embedded in optimal cutting temperature compound (Sakura, Finetek, Torrance, CA, USA) then snap frozen in liquid nitrogen. Lumbar spinal cords were sectioned into 16 µm thick transverse and coronal sections and dry mounted onto Superfrost Plus slides (Menzel-Glaser, Braunschweig, Germany) for quantitation of astrocytes and microglia as detailed below.

### Estimation of astrocytes and microglia

For estimation of astrocytes and microglia within the lumbar spinal cord, sections were rehydrated in PBS (pH 7.4) then blocked in PBS containing 3% bovine serum albumin (BSA) for 1 hour at room temperature. Sections were incubated overnight at 4°C with the astrocyte (mouse anti-GFAP; 1:1000, BD Biosciences, San Diego, CA, USA) and microglia (rat anti-CD11b; 1:500, Abcam, Cambridge, MA, USA and rat anti-CD68; 1:500, Bio-Rad, CA, USA) markers. Sections were washed with PBS for 3 × 10 minutes prior to incubation overnight at 4°C with the Alexa secondary cocktail: Alexa Fluor 555 dye-conjugated goat anti-rat (1:1000, Invitrogen, Eugene, OR, USA) and Alexa Fluor 488 dye-conjugated goat anti-mouse (1:600, Invitrogen, Eugene, OR, USA) antibody. All primary and secondary antibodies were diluted in PBS (pH 7.4) containing 1% BSA. Sections were then washed for 3 × 5 minutes in PBS, then mounted with Prolong Gold Anti-Fade medium containing 4, 6-diamidino-2-phenylindole (DAPI; Invitrogen, Eugene, OR, USA). Quantification of GFAP, CD11b and CD68 was performed on ∼18 to 21 lumbar spinal cord sections spaced 320µm apart and expressed as the percentage immunoreactive area per section [17,20]. Quantification was within the second lumbar dorsal root ganglia (L2) to the fifth lumbar dorsal root ganglia (L5), selected with the aid of the mouse spinal cord atlas [21]. Staining procedures and image exposures were all standardized between genotypes and between sections. The mouse genotype was not made available to the researchers until the completion of the study.

### Statistical analysis

All analyses were performed using GraphPad Prism 8.4.2 (San Diego, CA, USA). For the results from quantitative real-time PCR measuring Ager expression, statistical difference between SOD1^G93A^, TDP43^Q331K^ and TDP43^WTxQ331K^ mice and its corresponding WT controls were determined using a two-tailed student *t*-test for each age. The statistical difference between SOD1^G93A^ and SOD1^G93A^ x RAGE^-/-^ mice for GFAP, CD11b and CD68 quantification, were determined using two-tailed student *t*-test. The statistical difference for survival analyses between SOD1^G93A^ and SOD1^G93A^ x RAGE^-/-^ mice were analysed using log rank (Mantel-Cox) test. The statistical difference between SOD1^G93A^ and SOD1^G93A^ x RAGE^-/-^ mice for body weight, hind-limb grip strength, rota-rod and disease progression was analysed using a two-way ANOVA and a *post hoc* Fisher’s least significant difference test for each time point. For the results from quantitative real-time PCR, statistical difference between WT, RAGE^-/-^, SOD1^G93A^ and SOD1^G93A^ x RAGE^-/-^ in spinal cord and tibialis anterior muscle was determined using a one-way ANOVA and *post hoc* Tukey’s test. All data are presented as mean ± SEM and the differences were considered significant when *P* < 0.05.

## RESULTS

### Rage mRNA level is up-regulated progressively with disease severity in the spinal cord and tibialis anterior muscle of SOD1^G93A^, TDP43^Q331K^ and TDP43^WTxQ331K^ mice

We initially examined the mRNA expression of RAGE in the lumbar spinal cord and tibialis anterior (TA) muscle during key disease stages in SOD1^G93A^, TDP43^Q331K^ and TDP43^WTxQ331K^ mice using quantitative real-time PCR. In the SOD1^G93A^ mice, *Ager* transcript was initially decreased by 0.33-fold at onset, while it increased by 2.5-fold at the end-stage of disease in the lumbar spinal cord, when compared with WT mice (**Figure 1A**). In addition to the SOD1^G93A^ mice, *Ager* transcript was also increased in the lumbar spinal cord of TDP43^Q331K^ mice by 1.6-fold at 16 months of age (**Figure 1B**) and TDP43^WTxQ331K^ mice by 8.3-fold at 5 weeks of age (**Figure 1C**). *Ager* transcript in TA muscle of SOD1^G93A^ mice progressively increased by 1.6- and 5.6-fold at mid-symptomatic and end-stage of disease, respectively, when compared to WT mice (**Figure 1D**). Moreover, RAGE also displayed a marked increase in mRNA levels by 1.5-fold in TA muscle of TDP43^Q331K^ mice (**Figure 1E**) and 6.7-fold in TA muscle of TDP43^WTxQ331K^ mice (**Figure 1F**). Taken together, these results indicate that increased RAGE activation occurs in the lumbar spinal cord and TA muscle across multiple preclinical mouse models of ALS, suggesting a common pathway to glial activation and neuroinflammation, and ultimately disease progression in these models.

**Figure 1.**
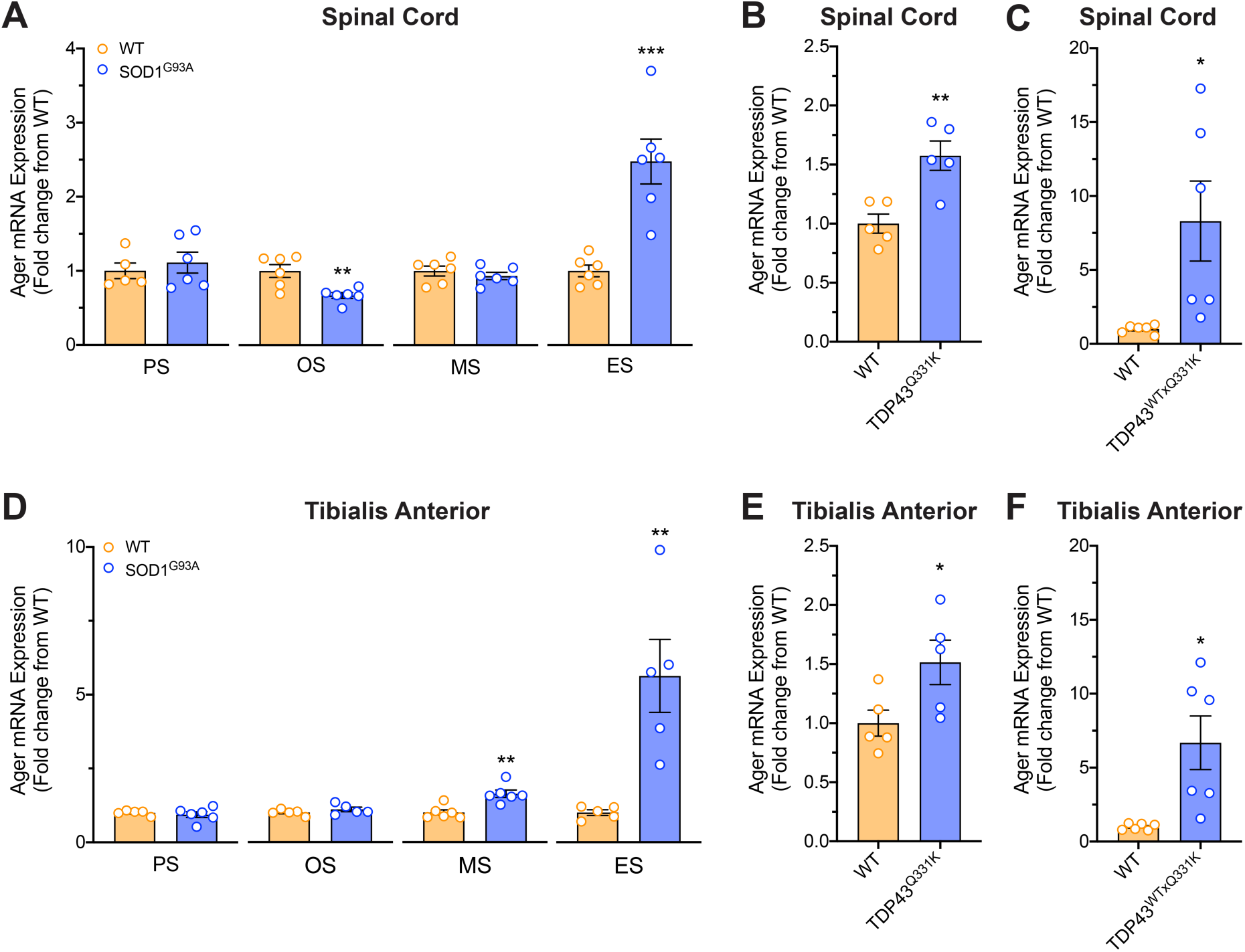
Expression of RAGE in the spinal cord and tibialis anterior muscle of SOD1^G93A^, TDP43^Q331K^ and TDP43^WTxQ331K^ mice. Panel **A** show the mRNA expression profile of *Ager* in the lumbar spinal cord of SOD1^G93A^ mice relative to wild-type (WT) mice at pre-symptomatic (PS; P30), onset (OS; P70), mid-symptomatic (MS; P130) and end-stage (ES; P155 - 175) of disease. **B** and **C** shows the mRNA expression profile of *Ager* in the lumbar spinal cord of TDP43^Q331K^ and TDP43^WTxQ331K^ mice relative to WT mice respectively at 16 months and 5 weeks of age. Panel **D** show the mRNA expression profile of *Ager* in the tibialis anterior muscle of SOD1^G93A^ mice relative to WT mice at PS, OS, MS and ES of disease. **E** and **F** shows the mRNA expression profile of *Ager* in the tibialis anterios muscle of TDP43^Q331K^ and TDP43^WTxQ331K^ mice relative to WT mice respectively at 16 months and 5 weeks of age. Data are expressed as means ± SEM (*n* = 5 - 6 mice/ group, * *P* < 0.05, ** *P* < 0.01, *** *P* < 0.001, two-tailed student *t*-test for each stage).

### Genetic deletion of RAGE improves hind-limb grip strength, rota-rod performance, extends survival and slows disease progression in SOD1^G93A^ mice

We next assessed whether enhanced RAGE signalling contributes to disease pathogenesis in ALS mice by generating SOD1^G93A^ mice entirely lacking RAGE (SOD1^G93A^ x RAGE^-/-^). SOD1^G93A^ x RAGE^-/-^ mice showed significantly extended survival when compared to SOD1^G93A^ mice (median end stage of disease, SOD1^G93A^ = 162 days and SOD1^G93A^ x RAGE^-/-^ = 179 days; **Figure 2A**). The body weight of SOD1^G93A^ and SOD1^G93A^ x RAGE^-/-^ mice reached their maximum at 119 and 112 days of age respectively; however, there was no difference in body weight loss between SOD1^G93A^ and SOD1^G93A^ x RAGE^-/-^ mice (**Figure 2B**). Motor deficits were also assessed in these animals using hind-limb grip strength and rota-rod performance. Concomitant with enhanced survival, there were significant improvements in hind-limb grip strength of SOD1^G93A^ x RAGE^-/-^ mice when compared to SOD1^G93A^ mice at 112, 119, 126, 133 and 140 days of age (**Figure 2C**). There were also significant improvements in rota-rod performance in SOD1^G93A^ x RAGE^-/-^ mice when compared to SOD1^G93A^ mice at 98, 105, 112, 119, 126, 133, 140, 147, 154 and 161 days of age (**Figure 2D**). Further to the improvement in motor function, overall disease progression determined by the age at which the maximal grip strength declined by 25, 50, 75 and 100%, was significantly delayed in SOD1^G93A^ x RAGE^-/-^ mice (**Figure 2E**).

**Figure 2.**
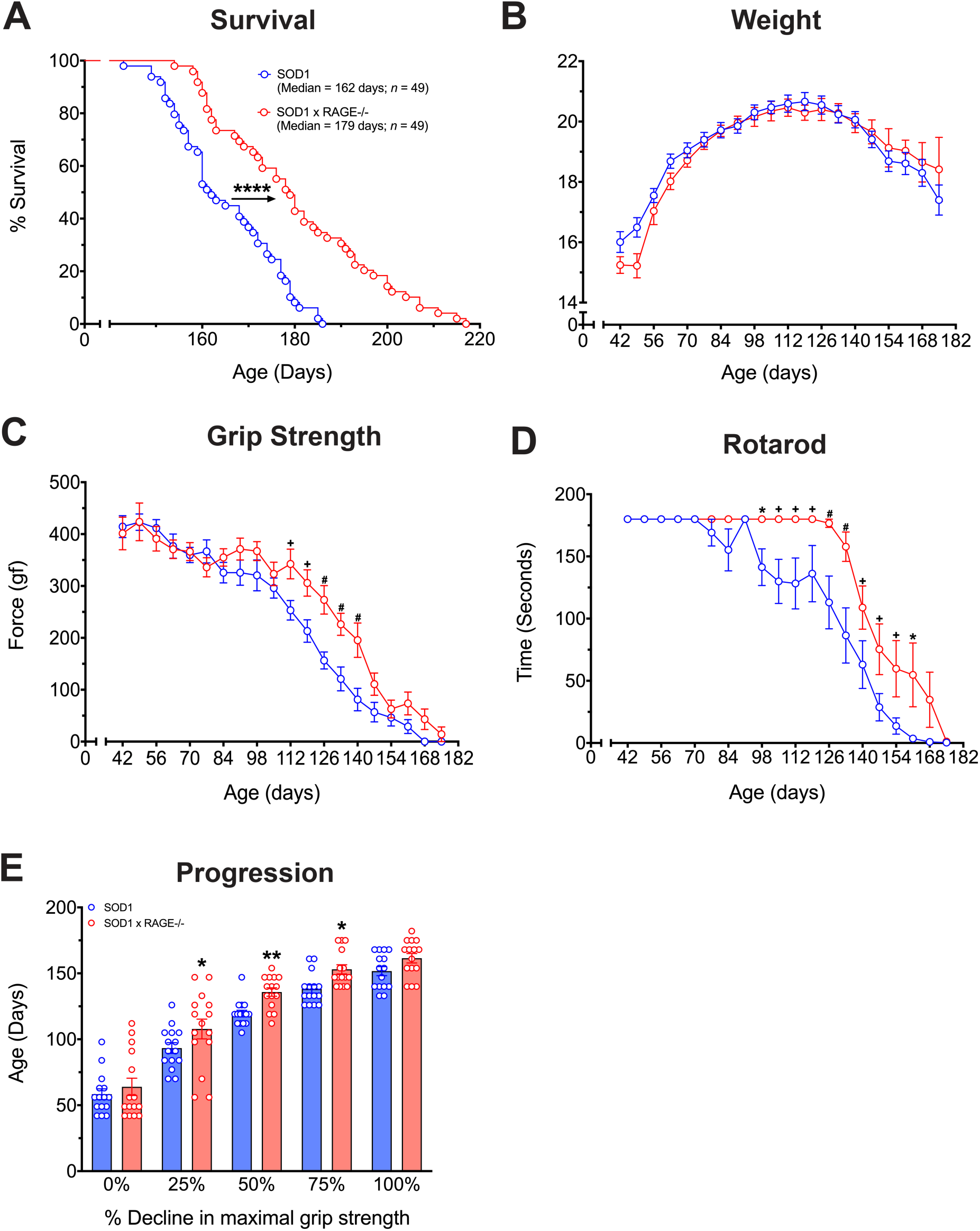
Genetic deletion of RAGE in SOD1^G93A^ mice extends survival time, improves motor performance and slows disease progression. **A** shows a Kaplan-Meier plot of ages (in days) in which SOD1^G93A^ mice with normal (RAGE^+/+^, blue line) or fully deleted (RAGE^-/-^, red line) RAGE reached end-stage of disease (complete hind-limb paralysis and an inability to right itself once placed on its back). SOD1^G93A^ x RAGE^-/-^ mice show a significant extension in survival time relative to SOD1^G93A^ mice (*n* = 49, **** *P* < 0.0001, log-rank test). **B** shows no difference in body weight between SOD1^G93A^ (blue line) and SOD1^G93A^ x RAGE^-/-^ mice (red line; *n* = 15, *P* = 0.3284, two-way ANOVA). **C** shows significant improvements in hind-limb grip strength of SOD1^G93A^ x RAGE^-/-^ mice (red line) when compared to SOD1^G93A^ mice (blue line) at 112, 119, 126, 133 and 140 days of age (*n* = 15, + *P* < 0.01, # *P* < 0.001; two-way ANOVA with *post hoc* Fisher’s LSD test at each age). There were also significant improvements in rota-rod performance in SOD1^G93A^ x RAGE^-/-^ (red line) when compared to SOD1^G93A^ mice (blue line) at 98, 105, 112, 119, 126, 133, 140, 147, 154 and 161 days of age (**D**; *n* = 15, * *P* < 0.05, + *P* < 0.01, # *P* < 0.001; two-way ANOVA with *post hoc* Fisher’s LSD test at each age). **E** shows significantly slowed disease progression (determined by age at which maximal grip strength decline at 25, 50, 75 and 100%) in SOD1^G93A^ x RAGE^-/-^ mice (red line) when compared with SOD1^G93A^ mice (blue line) at 25, 50 and 75% decline in maximal grip strength (*n* = 15, * *P* < 0.05, ** *P* < 0.01; two-way ANOVA with *post hoc* Fisher’s LSD test at each quartile). Data are expressed as mean ± SEM.

### Genetic deletion of RAGE reduces microglia and astrocyte accumulation in the spinal cord of SOD1^G93A^ mice

Microglia and astrocytes have been shown to play a key role in the pathogenesis of ALS [22-25]. Numerous studies have also linked RAGE signalling with modulation of gliosis [26-29,11], hence we examined glial markers in SOD1^G93A^ and SOD1^G93A^ x RAGE^-/-^ mice. We first investigated whether genetic deletion of RAGE in SOD1^G93A^ mice induced any effect on microglia and astrocytes expression markers in the lumbar spinal cord. mRNA expression levels of *Itgam, Cd68* and *Aif1* (markers of both resident microglia and infiltrating monocyte/macrophages) were measured in the lumbar spinal cord of SOD1^G93A^ and SOD1^G93A^ x RAGE^-/-^ mice at mid-symptomatic stage of disease progression using quantitative real-time PCR. *Itgam* transcripts were unaltered in SOD1^G93A^ x RAGE^-/-^ mice when compared to SOD1^G93A^ mice (**Figure 3A**). However, *Cd68* and *Aif1* transcripts were significantly reduced in SOD1^G93A^ x RAGE^-/-^ mice when compared to SOD1^G93A^ mice (**Figure 3B** and **3C**). Microglia were also examined using immunofluorescence (**Figure 3D**). No change in immunoreactive area of CD11b-positive microglia in the lumbar spinal cord of SOD1^G93A^ or SOD1^G93A^ x RAGE^-/-^ mice were found at mid-symptomatic stage of disease (**Figure 3E**). By contrast, a ∼50% reduction in the immunoreactive area of CD68-positive microglia was evident in the lumbar spinal cord of SOD1^G93A^ x RAGE^-/-^ mice at mid-symptomatic stage when compared to SOD1^G93A^ mice (**Figure 3F**). Next, we investigated the mRNA expression levels of *Gfap* (marker of astrocytes) in the lumbar spinal cord of SOD1^G93A^ and SOD1^G93A^ x RAGE^-/-^ mice at mid-symptomatic stage of disease. Similar to microglial markers, *Gfap* transcripts were also reduced in SOD1^G93A^ x RAGE^-/-^ mice when compared to SOD1^G93A^ mice (**Figure 3G**). Surprisingly, despite this transcript decrease, the immunoreactive area of GFAP-positive astrocytes did not change between SOD1^G93A^ and SOD1^G93A^ x RAGE^-/-^ mice (**Figure 3H** and **3I**).

**Figure 3.**
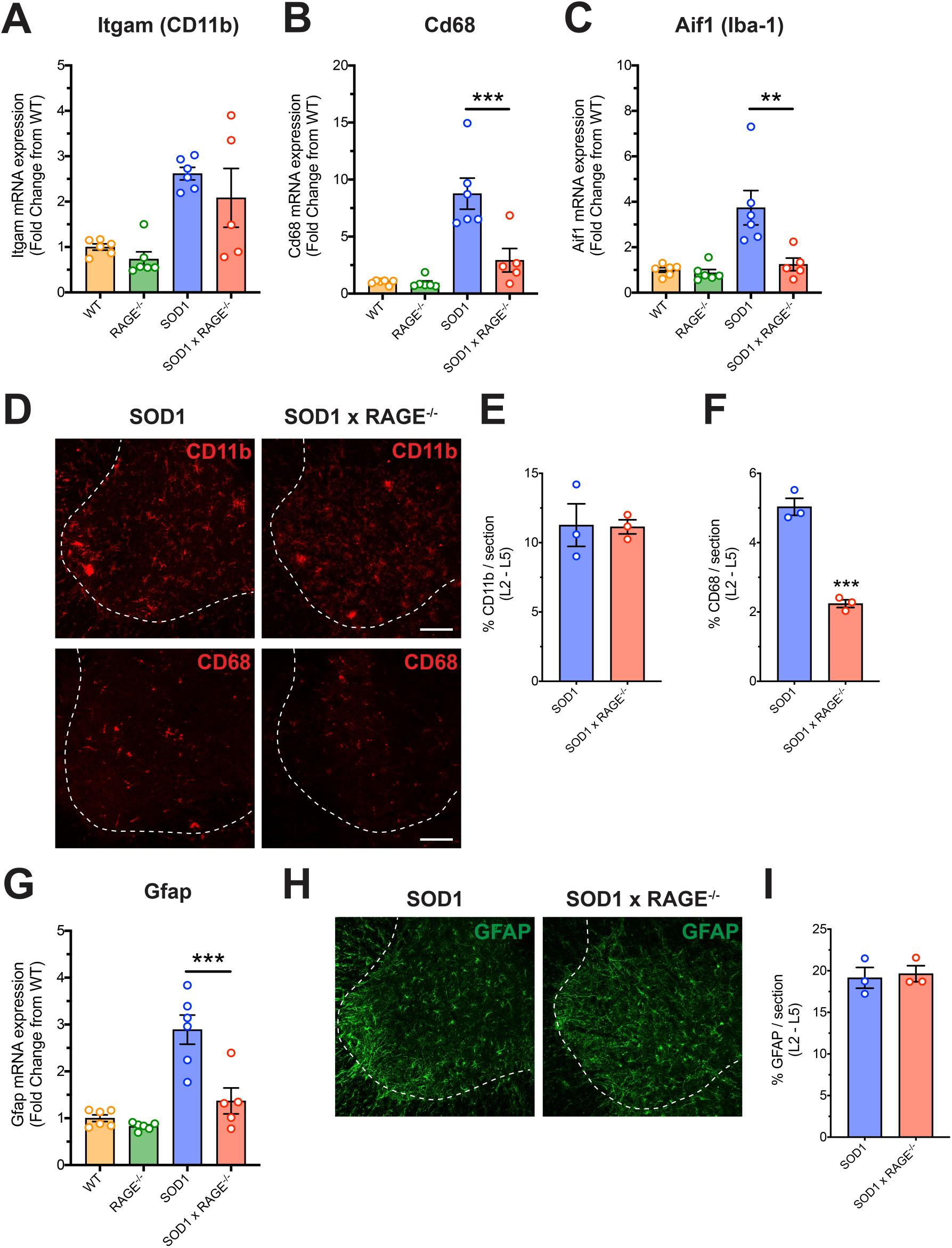
Genetic deletion of RAGE reduces glial markers in the spinal cord of SOD1^G93A^ mice. Major inflammatory cells in the spinal cord (microglia/monocytes and astrocytes) in WT, SOD1^G93A^, RAGE^-/-^ and SOD1^G93A^ x RAGE^-/-^ mice were investigated at mid-symptomatic stage of disease (130 days) using quantitative PCR and immunohistochemistry. **A** shows no difference in *Itgam* transcript between SOD1^G93A^ and SOD1^G93A^ x RAGE^-/-^ mice (*n* = 5 - 6, *P* = 0.6191, one-way ANOVA with *post hoc* Tukey’s test). **B** and **C** shows reduction in *Cd68* and *Aif1* mRNA transcript levels in SOD1^G93A^ x RAGE^-/-^ mice when compared to SOD1^G93A^ mice (*n* = 5 - 6, ** *P* < 0.01, *** P < 0.001, one-way ANOVA with *post hoc* Tukey’s test). **D** shows representative images of CD11b and CD68-positive microglia/macrophages in the lumbar spinal cord of SOD1^G93A^ and SOD1^G93A^ x RAGE^-/-^ mice at 130 days of age. White dashed line shows the outline of the ventral horn. Scale bar = 100 μm. **E** shows no change in immunoreactive area of CD11b-positive microglia expression in the lumbar spinal cord of SOD1^G93A^ or SOD1^G93A^ x RAGE^-/-^ mice (*n* = 3, *P* = 0.9416; Student’s t-test). A reduction in the immunoreactive area of CD68 positive microglia was evident in the lumbar spinal cord of SOD1^G93A^ x RAGE^-/-^ mice when compared to SOD1^G93A^ mice (**F**, *n* = 3, *** *P* < 0.001, Student’s t-test). **G** shows a decrease in *Gfap* mRNA transcript levels in SOD1^G93A^ x RAGE^-/-^ mice when compared to SOD1^G93A^ mice (*n* = 5 - 6, *\*\*\* P* < 0.001, Student’s t-test). **H** shows representative images of GFAP-positive astrocytes in the lumbar spinal cord of SOD1^G93A^ and SOD1^G93A^ x RAGE^-/-^ mice at 130 days of age. White dashed line shows the outline of the ventral horn. Scale bar = 100 μm. **I** show no change in the immunoreactive area of GFAP-positive astrocytes in the lumbar spinal cord of SOD1^G93A^ or SOD1^G93A^ x RAGE^-/-^ mice (*n* = 3, *P* = 0.7701, Student’s t-test). Data are presented as mean ± SEM.

### SOD1^G93A^ mice lacking RAGE have decreased spinal cord levels of key inflammatory markers

Activation of RAGE signalling induces synthesis of cytokines and initiates oxidative stress to propagate inflammatory processes. RAGE has been shown to induce cytokine expression specifically in microglia [27,11]. Importantly, pro-inflammatory cytokines such as TNFα and IL-1β are thought to propagate disease progression in ALS in connection with the innate immune system [30]. Hence, we investigated whether genetic deletion of RAGE in SOD1^G93A^ mice altered the expression of TNFα and IL-1β and several major components of the innate immune system including the complement system (C1q, C3 and C5aR1), toll-like receptor (HMGB1 and TLR4), inflammasome (NLRP3 and ASC) and oxidative stress (NOX2) in the lumbar spinal cord. mRNA expression of all the transcripts were measured in SOD1^G93A^ and SOD1^G93A^ x RAGE^-/-^ mice at mid-symptomatic stage of disease progression by quantitative real-time PCR. *Tnf* and *Il1b* transcripts were significantly reduced in SOD1^G93A^ x RAGE^-/-^ mice by 0.55- and 0.61-fold when compared to SOD1^G93A^ mice respectively (**Figure 4A** and **4B**). A significant reduction in transcripts associated with the complement system (*C1qb, C3* and *C5ar1*) was observed in SOD1^G93A^ x RAGE^-/-^ mice. Specifically, *C1qb, C3* and *C5ar1* mRNA expression was decreased in the lumbar spinal cord of SOD1^G93A^ x RAGE^-/-^ mice by 0.39-, 0.5- and 0.46-fold when compared to SOD1^G93A^ mice (**Figure 4C, 4D** and **4E**). *Hmgb1* and *Tlr4* transcripts were also reduced by 0.45- and 0.34-fold in the SOD1^G93A^ x RAGE^-/-^ mice respectively (**Figure 4F** and **4G**). Inflammasome component *Nlrp3* transcripts were significantly decreased in the lumbar spinal cord of SOD1^G93A^ x RAGE^-/-^ mice by 0.4-fold at mid-symptomatic stage when compared to SOD1^G93A^ mice (**Figure 4H**), while *Pycard* mRNA expression showed a decrease of 0.37-fold in SOD1^G93A^ x RAGE^-/-^ mice (**Figure 4I**). Lastly, *Cybb* transcripts (an oxidative stress marker) were reduced by 0.6-fold in the lumbar spinal cord of SOD1^G93A^ x RAGE^-/-^ mice (**Figure 4J**). Taken together these results suggest that genetic ablation of RAGE reduces major pro-inflammatory factors in the spinal cord of SOD1^G93A^ mice.

**Figure 4.**
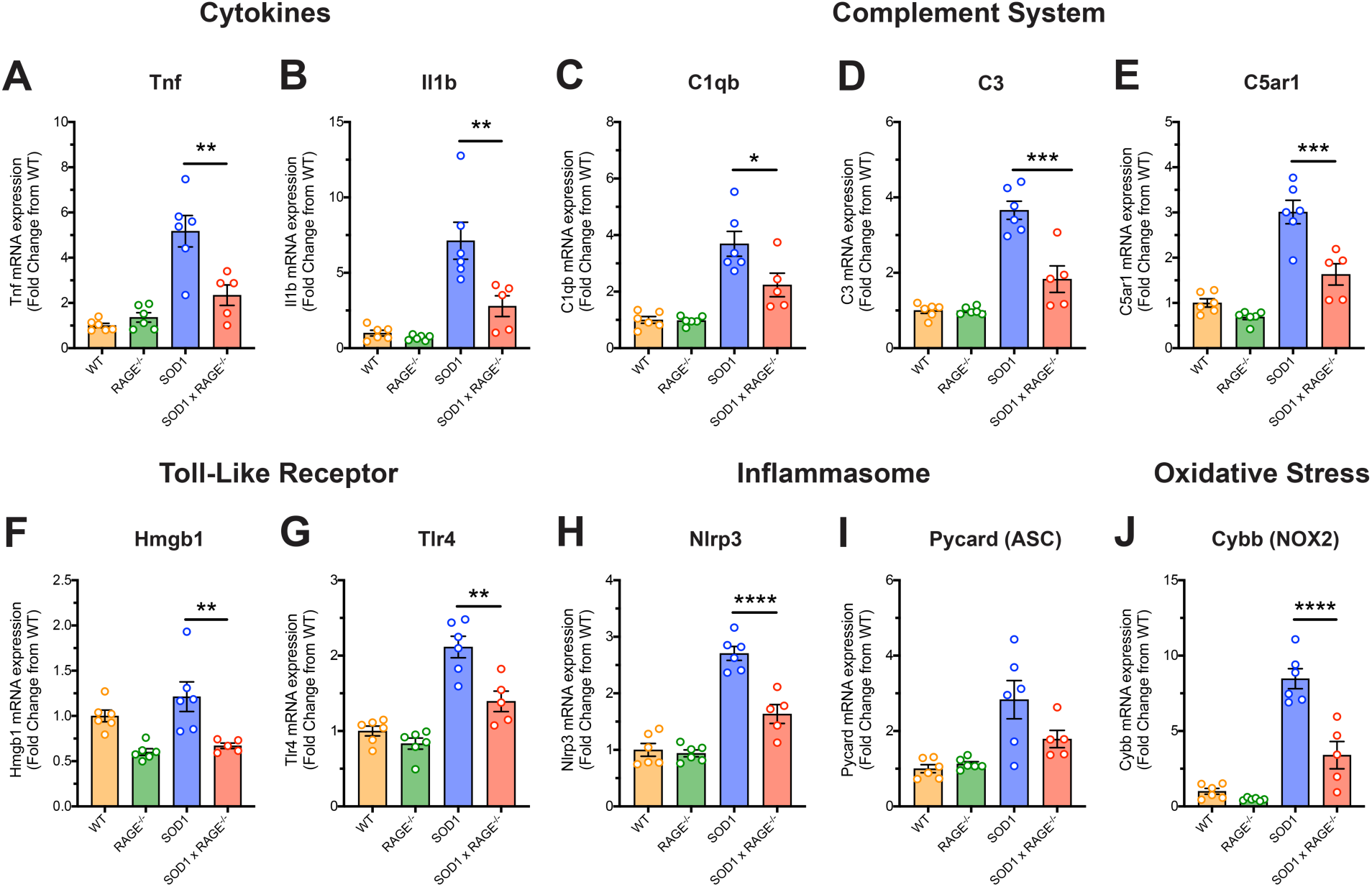
SOD1^G93A^ mice lacking RAGE have decreased spinal cord levels of key inflammatory markers. Pro-inflammatory cytokines (TNFα and IL-1β), complement components (C1q, C3 and C5aR1), toll-like receptor pathway components (HMGB1 and TLR4), inflammasome components (NLRP3 and ASC) and an oxidative stress marker (NOX2) in the lumbar spinal cord of WT, SOD1^G93A^, RAGE^-/-^ and SOD1^G93A^ x RAGE^-/-^ mice were investigated at mid-symptomatic stage of disease (130 days) using quantitative PCR. **A** and **B** shows decreased mRNA levels of pro-inflammatory cytokines *Tnf* and *Il1b* in SOD1^G93A^ x RAGE^-/-^ mice when compared to SOD1^G93A^ mice (*n* = 5 - 6, ** *P* < 0.01, one-way ANOVA with *post hoc* Tukey’s test). There was significant reduction in transcripts associated with the complement system (*C1qb, C3* and *C5ar1*) in the SOD1^G93A^ x RAGE^-/-^ mice when compared to SOD1^G93A^ mice (**C** - **E**, *n* = 5 - 6, * *P* < 0.05, *** *P* < 0.001, one-way ANOVA with *post hoc* Tukey’s test). **F** and **G** shows reduction in *Hmgb1* and *Tlr4* mRNA transcript levels in the SOD1^G93A^ x RAGE^-/-^ mice when compared to SOD1^G93A^ mice (*n* = 5 - 6, ** *P* < 0.01 one-way ANOVA with *post hoc* Tukey’s test). Inflammasome component *Nlrp3* transcript was significantly decreased in the SOD1^G93A^ x RAGE^-/-^ mice when compared to SOD1^G93A^ mice (**H**, *n* = 5 - 6, **** *P* < 0.0001, one-way ANOVA with *post hoc* Tukey’s test), while *Pycard* mRNA expression showed no difference between to SOD1^G93A^ and SOD1^G93A^ x RAGE^-/-^ mice (**I**, *n* = 5 - 6, *P* = 0.0966, one-way ANOVA with *post hoc* Tukey’s test). **J** shows a reduction in *Cybb* transcript levels in SOD1^G93A^ x RAGE^-/-^ when compared to SOD1^G93A^ mice (*n* = 5 - 6, **** *P* < 0.0001, one-way ANOVA with *post hoc* Tukey’s test). Data are presented as mean ± SEM.

### Genetic deletion of RAGE reduces denervation markers, but not macrophage or cytokine markers in the tibialis anterior muscle of SOD1^G93A^ mice

As genetic deletion of RAGE showed a marked reduction in microglial and pro-inflammatory markers in the spinal cord of SOD1^G93A^ mice, we next investigated whether peripheral macrophage infiltration and pro-inflammatory cytokine levels in the TA muscle were also impacted. mRNA expression levels of monocytes/macrophage markers *Itgam* and *Aif1* and pro-inflammatory cytokines *Tnf* and *Il1b* were therefore measured in the TA muscle of SOD1^G93A^ and SOD1^G93A^ x RAGE^-/-^ mice at mid-symptomatic stage using quantitative real-time PCR. Interestingly, we showed no change in these transcript levels in SOD1^G93A^ x RAGE^-/-^ mice when compared to SOD1^G93A^ mice (**Figure 5A - 5D**). As we identified a significant improvement in muscle function in SOD1^G93A^ x RAGE^-/-^ mice at mid-symptomatic age (**Figure 2C** and **2D**), we also examined denervation markers (Musk, Chrna1, Cpla2, Ncam1 and Runx1) using quantitative real-time PCR. We identified a significant reduction in the mRNA expression of *Musk* (0.48-fold; **Figure 6A**), *Chrna1* (0.54-fold; **Figure 6B**), *Cpla2* (0.45-fold; **Figure 6C**), Ncam1 (0.62-fold; **Figure 6D**) and *Runx1* (0.6-fold; **Figure 6E**) at mid-symptomatic stage in TA muscle of SOD1^G93A^ x RAGE^-/-^ mice when compared to SOD1^G93A^ mice. This suggests that while RAGE deficiency did not impact on inflammation markers, denervation was still reduced in affected SOD1^G93A^ muscles.

**Figure 5.**
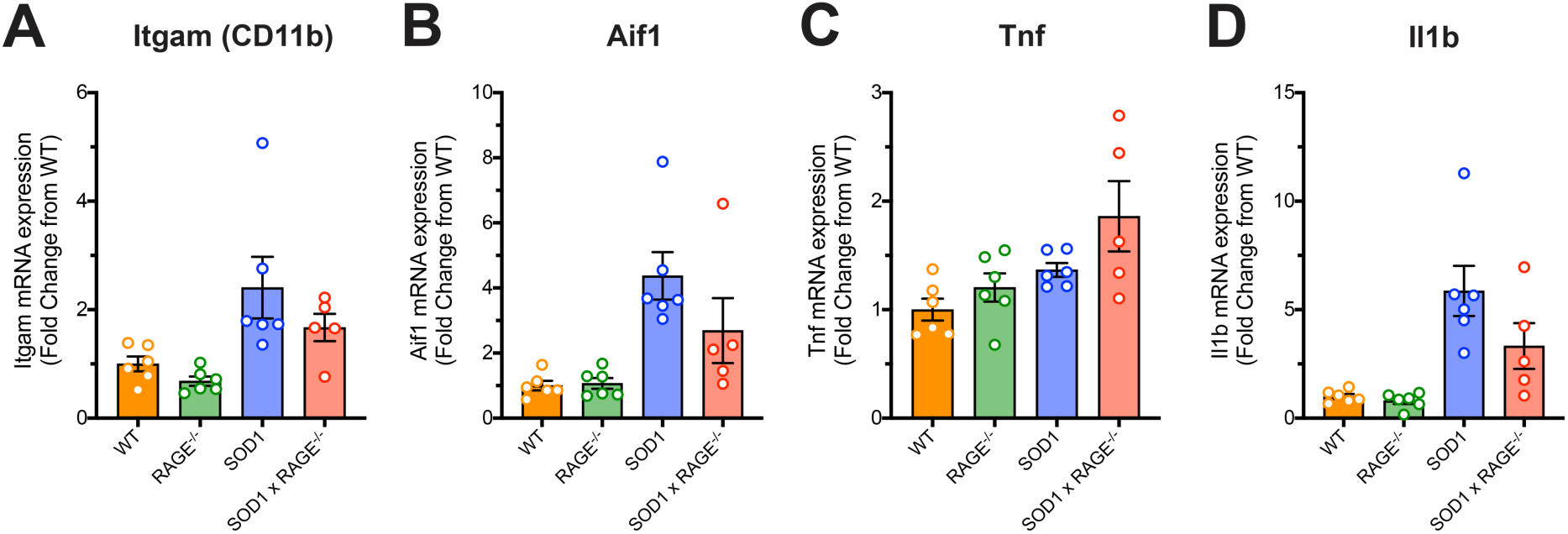
Lack of RAGE signaling in SOD1^G93A^ mice does not alter macrophage and pro-inflammatory cytokines in tibialis anterior muscle. Macrophage markers (*Itgam* and *Aif1*) and pro-inflammatory cytokines (*Tnf* and *Il1b*) in tibialis anterior muscle of WT, RAGE^-/-^, SOD1^G93A^ and SOD1^G93A^ x RAGE^-/-^ mice were investigated at mid-symptomatic stage (130 days) using quantitative PCR. Panel **A** to **D** shows no change in macrophage (*Itgam* and *Aif1*) and pro-inflammatory cytokine (*Tnf* and *Il1b*) mRNA transcript levels between SOD1^G93A^ and SOD1^G93A^ x RAGE^-/-^ mice (*n* = 5 - 6, Itgam: *P* = 0.4292, Aif1: *P* = 0.2301, Tnf: *P* = 0.2082, Il1b: *P* = 0.1349, one-way ANOVA with *post hoc* Tukey’s test). Data are presented as mean ± SEM.

**Figure 6.**
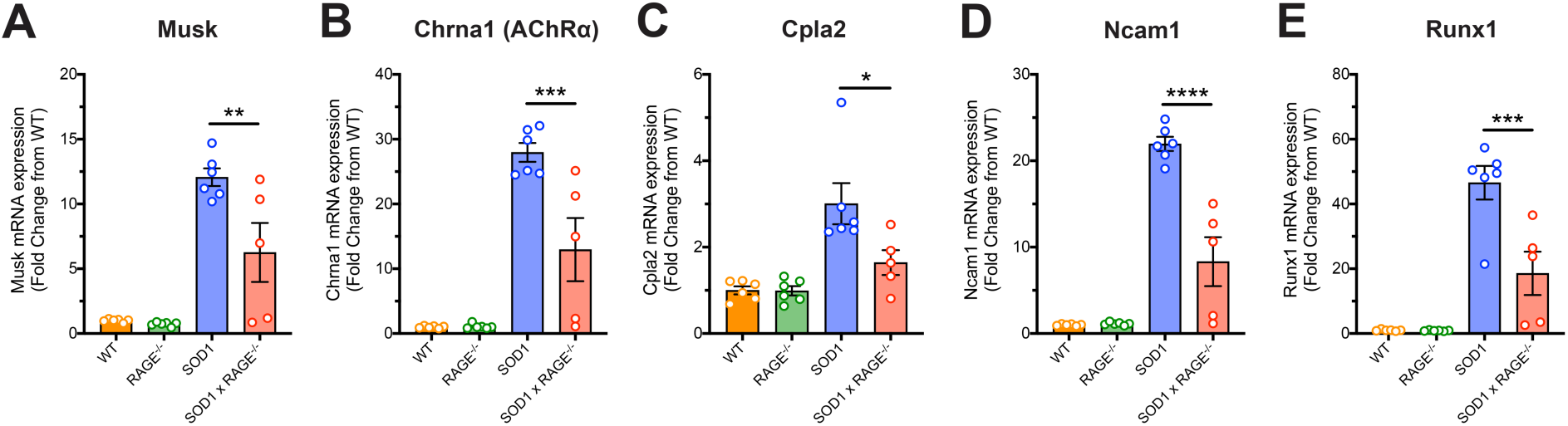
Genetic deletion of RAGE reduces denervation markers in the tibialis anterior muscle of SOD1^G93A^ mice. Denervation markers (*Musk, Chrna1, Cpla2, Ncam1* and *Runx1*) in tibialis anterior muscle of WT, Rage^-/-^, SOD1^G93A^ and SOD1^G93A^ x RAGE^-/-^ mice were investigated at mid-symptomatic stage (130 days) using quantitative PCR. Panel **A** to **E** shows a significant reduction in all denervation markers in SOD1^G93A^ x RAGE^-/-^ mice when compared to SOD1^G93A^ mice (*n* = 5 – 6, * P < 0.05, ** P < 0.01, *** P < 0.001, **** P < 0.0001, one-way ANOVA with *post hoc* Tukey’s test). Data are presented as mean ± SEM.

## DISCUSSION

The major findings of the current study are that RAGE is progressively upregulated in the spinal cord and skeletal muscle of multiple preclinical ALS mouse models, and that lack of RAGE signalling in SOD1^G93A^ mice exerts a protective effect on survival and motor function. This protection was associated with a reduction in widespread inflammation and oxidative stress markers in the spinal cord, and a decrease in denervation markers in the skeletal muscle.

It is now well documented that RAGE and its pro-inflammatory ligands are potentially involved in ALS, with evidence from both human patients and rodent models [6,10,11,13,20,19,12]. The present study further adds to this knowledge demonstrating alteration of RAGE mRNA expression in the spinal cord and skeletal muscle of multiple preclinical transgenic ALS mouse models including SOD1^G93A^, TDP43^Q331K^ and TDP43^WTxQ331K^ mice, suggestive of a common activation pathway for RAGE upregulation in these models, that is independent to the underlying genetic driver of the disease. Our results support prior studies in SOD1^G93A^ mice that highlighted RAGE activation in the spinal cord [10,12], and provide the first evidence demonstrating RAGE upregulation also occurs in the skeletal muscle. These data suggest that increased RAGE expression in these degenerating tissues occurs in response to motor neuron death and muscle denervation. Our findings also indicate that RAGE activation could be a common mechanism underlying most forms of ALS.

RAGE is a pro-inflammatory pattern recognition receptor expressed on microglia and astrocytes that plays an important role in inflammation, oxidative stress and cellular dysfunction in a number of neurodegenerative diseases including Alzheimer’s disease, Parkinson’s disease and ALS [4]. RAGE binds to danger associated molecular pattern (DAMP) proteins that are released from damaged or dying neurons and its interaction triggers multiple intracellular signaling pathways which promotes the production of pro-inflammatory molecules and reactive oxygen species, leading to oxidative stress and further neuronal loss [7,8]. In support of this, the present study demonstrated that homozygote deletion of RAGE in SOD1^G93A^ mice significantly reduced pathology by extending survival time, improving motor performance in hind-limb grip strength and rota-rod testing, and slowing disease progression. In addition to the significant improvement in muscle function we also observed a decrease in denervation markers in skeletal muscle. Our results therefore demonstrate that RAGE activation in SOD1^G93A^ mice actively contributes to spinal cord and skeletal muscle pathogenesis. This is consistent with previous studies where pharmacological inhibition of RAGE signaling using a soluble RAGE form that acts as a RAGE decoy, exerted beneficial effects on survival and disease progression in SOD1^G93A^ mice [10,12]. Inhibition of RAGE signaling has also been neuroprotective in a number of other neurodegenerative diseases [31,32,29,33]. Our data corroborates the premise that enhanced RAGE signaling plays a key role in accelerating motor neuron loss and neuromuscular junction denervation, and ultimately accelerating ALS progression.

In line with the beneficial effect on SOD1^G93A^ pathology from genetic deletion of RAGE, our studies also demonstrated a reduction in microglia (both mRNA and protein markers), and astrocytes (mRNA markers only) in the spinal cord. This fits with current hypotheses regarding the neuropathogenic mechanisms of enhanced RAGE activation, as numerous studies have linked RAGE signalling with modulation of gliosis [26-29,11]. Activation of RAGE triggers multiple signalling pathways that leads to the production of pro-inflammatory mediators including cytokines, nitric oxide and reactive oxygen species [34,8]. Many of these factors have been shown to be neurotoxic to motor neurons, and contribute to ALS pathology [35-39]. Therefore, we hypothesised that enhanced RAGE signalling in SOD1^G93A^ animals could drive neuroinflammatory responses that accelerates motor neuron loss. The present study clearly demonstrated a marked reduction in overall inflammatory markers in the spinal cord including decreases in pro-inflammatory cytokines TNF*α* and IL-1β, the RAGE ligand and danger-associated molecular pattern molecule HMGB1, innate immune receptors TLR4 and C5aR1, inflammasome component NLRP3, and an oxidative stress marker. This is consistent with many prior studies demonstrating these components play pathogenic roles in ALS disease progression [35,36,40-43,20,19,37,44]. Interestingly, genetic deletion of RAGE also resulted in a down-regulation of the complement components C1qb and C3. Previous studies have shown that RAGE can directly bind to C1q to enhance C1q-mediated phagocytosis and that RAGE signalling through p38MAPK and NF-*κ*B can induce C3 production [45,46]. Hence, it is plausible that the absence of RAGE signalling could reduce the removal of motor neurons via opsonisation and removal of neuromuscular synapses via phagocytosis. Furthermore, the observed reduction in GFAP and C3 mRNA levels, could indicate that deletion of RAGE resulted in a reduction of neurotoxic A1 astrocyte accumulation, of which C3 is a major cell marker [24].

Although genetic deletion of RAGE resulted in a reduction of overall inflammation in the spinal cord of SOD1^G93A^ mice, surprisingly, macrophage and pro-inflammatory cytokine markers within the skeletal muscle were not altered. This is contradictory to a prior study demonstrating ablation of RAGE resulted in reductions in lympho-mononuclear cell infiltrates and pro-inflammatory factors in damaged muscle tissue of mice with Duchenne muscular dystrophy [47]. One explanation for this difference could be due to the difference in pathological mechanisms leading to muscular dysfunction, where symptoms of muscle weakness in ALS originates from motor neuron loss in the motor cortex and spinal cord, while muscle weakness in Duchenne muscular dystrophy originates from within the muscle itself with a complete absence of an essential component of the dystrophin-associated protein complex located at the sarcolemma [1,47]. Our results indicate that the effects of RAGE signalling on inflammation in the SOD1^G93A^ model is primarily driven centrally, rather than peripherally. This is further supported by our demonstration of reduced denervation markers in the skeletal muscle, despite observing no change inflammation markers in the same tissue samples. It is therefore likely that amelioration of spinal cord inflammation following the absence of RAGE in SOD1^G93A^ mice, led to decreased motor neuron loss, subsequently reducing skeletal muscle denervation markers, however, this hypothesis would require further experimental validation.

In summary, the present study demonstrated that RAGE is upregulated in the spinal cord and skeletal muscle of multiple preclinical ALS mouse models. The genetic deletion of RAGE in SOD1^G93A^ ALS mice significantly extended survival time, improved motor performance and slowed disease progression, associated with reduced spinal cord expression of key pro-inflammatory and oxidative stress molecules and skeletal muscle expression of denervation markers. These data therefore indicate that RAGE signalling plays a pathogenic role in the SOD1^G93A^ mouse model of ALS, and together with our expression data showing RAGE elevation in TDP43 ALS models, highlights RAGE as a potential immune therapeutic target for all forms of ALS.

## ACKNOWLEDGEMENTS

The authors would like to sincerely thank Kym French for the animal care and husbandry. We also thank Maryam Shayegh for her technical support with genotyping the mice, and A/Prof Simon Phipps for original supply of RAGE^-/-^ breeders used to establish our colony.

## DECLARATION

### Ethics approval

All experimental procedures were approved by the University of Queensland Animal Ethics Committee, and complied with the policies and regulations regarding animal experimentation and other ethical matters. They were conducted in accordance with the Queensland Government Animal Research Act 2001, associated Animal Care and Protection Regulations (2002 and 2008), and the Australian Code of Practice for the Care and Use of Animals for Scientific Purposes, 8^th^ Edition (National Health and Medical Research Council, 2013).

### Consent to participate

Not applicable

### Consent for publication

Not applicable

### Availability of data and material

Not applicable

### Code availability

Not applicable

### Author’s contributions

JDL and TMW conceived the project. JDL and TMW designed the study. JDL performed the experiments with the assistance from TSM and JNTF. All authors contributed to the analyses and interpretation of the data. JDL wrote the paper with contribution from TMW. All authors read and approved the final manuscript.

